# Centrosome-nuclear envelope tethering and microtubule motor-based pulling forces collaborate in centrosome positioning during mitotic entry

**DOI:** 10.1101/442368

**Authors:** Vincent Boudreau, Richard Chen, Alan Edwards, Muhammad Sulaimain, Paul S. Maddox

## Abstract

Centrosome positioning relative to the nucleus and cell shape is highly regulated across cell types, during cell migration and during spindle formation in cell division. Across most sexually reproducing animals, centrosomes are provided to the oocyte through fertilization and must be positioned properly to establish the zygotic mitotic spindle. How centrosomes are positioned in space and time through the concerted action of key mitotic entry biochemical regulators including Protein Phosphatase 2A (PP2A-B55/SUR-6), biophysical regulators including Dynein and the nuclear lamina is unclear. Here, we uncover a role for PP2A-B55/SUR-6 in regulating centrosome positioning. Mechanistically, PP2A-B55/SUR-6 regulates nuclear size prior to mitotic entry, in turn affecting nuclear envelope-based Dynein density and motor capacity. Using computational simulations, PP2A-B55/ SUR-6 regulation of nuclear size and nuclear envelope Dynein density were both predicted to be required for proper centrosome positioning. Conversely, compromising nuclear lamina integrity led to centrosome detachment from the nuclear envelope and migration defects. Removal of PP2A-B55/SUR-6 and the nuclear lamina simultaneously further disrupted centrosome positioning, leading to unseparated centrosome pairs dissociated from the nuclear envelope. Taken together, we propose a model in which centrosomes migrate and are positioned through the concerted action of nuclear envelope-based Dynein pulling forces and cen-trosome-nuclear envelope tethering.

## INTRODUCTION

Fertilization provides centrosomes to the developing embryo in most known systems. Associated to the sperm pronucleus, centrosomes must grow, separate and migrate in a coordinated fashion to facilitate female and male pronuclear meeting prior to mitosis. In doing so, centrosomes orchestrate the microtubule cytoskeleton in the embryo such that the mitotic spindle is properly positioned, ensuring polarity during asymmetric cell division. Simultaneously, as the zygote enters mitosis, collaboration between mitotic master regulators and microtubule-based biophysical regulators must be organized.

Mechanistically, centrosomes are positioned through the concerted action of cytoplasmic flow, cytoplasmic and cortical microtubule pulling forces, and microtubule pushing forces against the cortex (Garzon-Coral et al., 2016; Nazockdast et al., 2017). On the molecular scale, it has been well established that microtubule-based motors including Dynein (Gonczy et al., 1999; Robinson et al., 1999) and Eg5 (Blangy et al., 1995; Sawin et al., 1995) play important roles in centrosome migration at mitotic entry. Recently, a combination of experimental and theoretical approaches have been used to distinguish the roles of different pools of motors (i.e. nuclear and cortical), cortical flows (De Simone et al., 2016, 2017) and cytoplasmic drag (De Simone et al., 2018) in regulating centrosome positioning at the mesoscale. In the *C. elegans* zygote, Dynein-based microtubule pulling forces are produced at the nuclear envelope and at the cortex to ensure proper centrosome positioning (De Simone et al., 2016). Dynein regulators including LIS-1 are also required for centrosome separation in the embryo (Cockell et al., 2004), although their spatio-temporal contributions remain elusive. Despite our understanding of microtubule cyto-skeleton-based pulling forces, the contribution of key mitotic kinases and phosphatases to centrosome positioning remains unclear.

The net effect of molecular regulators is a biophysical mechanism required for positioning centrosomes during mitotic entry in the early embryo and the contributions of the nuclear envelope and its centrosome tethering components are as yet uncertain. Centrosomes are tethered to the nuclear lamina through the linker of nucleoskeleton and cytoskeleton (LINC) complex, which is composed of SUN proteins tethered to the nuclear lamina and KASH proteins tethered to the microtubule cytoskeleton (reviewed in Chang et al., 2015). In the *C. elegans* embryo, both the SUN protein SUN-1 and the KASH protein ZYG-12 are required for centrosome positioning (Malone et al., 2003). Centrosome-nuclear envelope tethering is thought to be achieved through Dynein as ZYG-12 directly binds Dynein’s light chain (Malone et al., 2003) and Dynein is required for tethering (Gonczy et al., 1999). Dynein-based microtubule pulling forces and the centro-some-nuclear envelope tethering machinery are therefore likely to be coordinated in centrosome positioning during mitotic entry.

Biochemically, mitotic entry is defined by the activities of master mitotic kinases and phosphatases that ensure faithful cell cycle progression (Krasinka et al., 2011; Mochida et al, 2016). As Cyclin Dependent Kinase 1 (CDK1) activity increases at mitotic entry, its main counteracting phosphatase Protein Phosphatase 2A (PP2A) is also regulated. In metazoans, PP2A mainly functions as a heterotrimeric enzyme using its B55/SUR-6 adaptor subunit for substrate recognition and is the main phosphatase responsible for dephosphorylating CDK1 substrates (Mayer-Jaekel et al., 1994; Castilho et al., 2009; Mochida et al., 2009). To identify potential converse functions of the CDK1 counteracting phosphatase PP2A-B55/SUR-6, we previously used a maternal effect genetic screen in the fly embryo (Mehsen et al., 2018) in which several components were identified as PP2A-B55/SUR-6 collaborators for proper cell cycle progression, including Lamin. The biology of *D. melanogaster* early development (e.g. the zygotic division) precluded centrosome tracking and therefore fundamental probing of the biophysical mechanisms at play, thus we examined the contributions of the genetic interaction between PP2A-B55/ SUR-6 and Lamin/LMN-1 to the first developmental mitosis in the *C. elegans* embryo (the zygote). We identify a novel role for PP2A-B55/SUR-6 in regulating centrosome positioning in the embryo through the regulation of nuclear envelope-tethered Dynein motor activity. We propose a simple model in which Dynein-based pulling forces collaborate with centrosome-nuclear envelope tethering to ensure proper centrosome positioning during mitotic entry.

## RESULTS

### PP2A-B55/SUR-6 and LMN-1 play critical roles in centrosome positioning in the C. elegans zygote

To identify potential roles for the uncovered genetic interactions in single embryonic mitoses, we depleted individual components by RNAi in the *C. elegans* embryo – a system in which late meiotic and the first mitotic events can be readily observed by light microscopy (<u>Video S1</u>) (Fig. 1A). Depleting *sur-6*, the PP2A adapter subunit, in worms expressing H2B:GFP and γ-tubulin:GFP (strain TH32) led to centrosomes that were tethered to the nuclear envelope and failed to separate from one another (<u>Video S2</u>) (Fig. 1B and F) while depleting *lmn-1* caused centrosomes to detach from nuclear envelopes and dramatically increase in distance between one another (<u>Video S3</u>) (Fig. 1D and G).

We hypothesized that PP2A-B55/SUR-6 may regulate centrosome migration through biochemical regulation of known centrosome separation components including Dynein. Partial RNAi-mediated depletion of the Dynein motor heavy chain *dhc-1* led to centrosome migration defects while centrosomes remained tethered to the nuclear envelope (<u>Video S4</u>) (Fig. 1C and F). Similarly, depleting the Dynein regulator *lis-1* led to centrosome migration phenotypes while centrosome tethering was maintained (<u>Video S5</u>) (Fig. S1B). Taken together, partial depletion of Dynein’s heavy chain or of its regulator *lis-1* phenocopied *sur-6* centrosome migration phenotypes, supporting a potential role for PP2A-B55/ SUR-6 in regulating centrosome positioning through Dynein activity.

**Figure 1.**
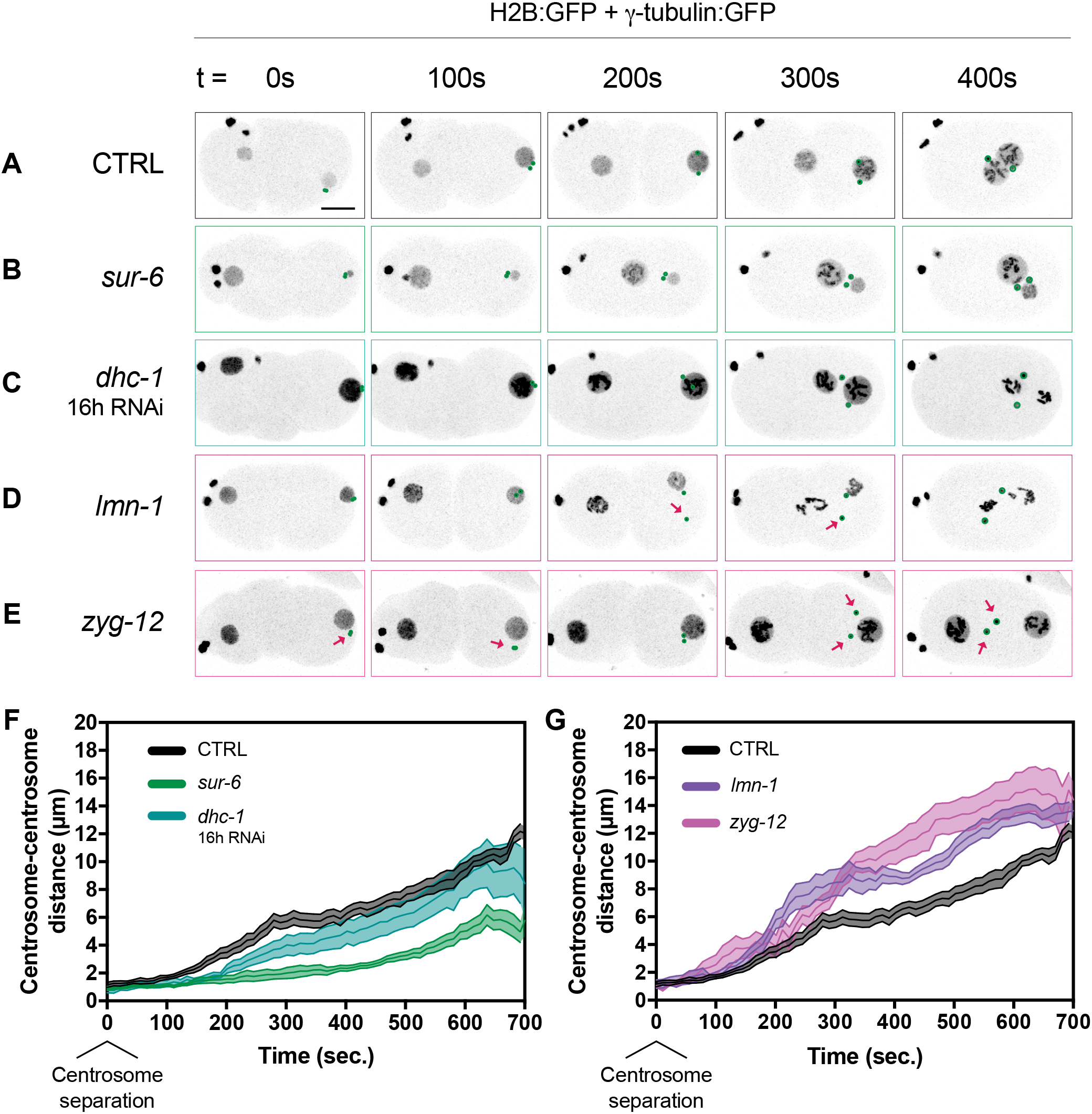
*sur*-6 and *lmn-1* RNAi affect centrosome separation akin to *dhc-1* and *zyg-12*. (A)(B)(C)(D)(E) Representative maximum intensity projections of embryonic confocal stacks. All RNAi treatments were performed with L4 stage worms expressing H2B:GFP and γ-tubulin:GFP for 24 hours via feeding unless otherwise noted. Circles highlight individual centrosomes and arrows denote detached centrosomes. Scale bar = 10 μm. (F)(G) Centrosome distance was measured in 2D using maximum intensity projections of confocal stacks. Error bars are SEM and are represented as shaded areas around means. n of embryos analyzed per condition > 15. Time t = 0 s corresponds to centrosome separation.

Given LMN-1’s role as the principal structural component of the nuclear lamina, we hypothesized that *lmn-1* cen-trosome migration phenotypes may be the result of failed centrosome-nuclear envelope cohesion. Consistent with previous reports, depletion of the outer nuclear envelope LINC complex component *zyg-12* led to the loss of centrosome-nuclear envelope cohesion and increased centrosome-centrosome distance (<u>Video S6</u>) (Fig. 1E and G). Similarly, depleting the inner nuclear envelope LINC complex component *sun-1* compromised centrosome-nu-clear envelope tethering (<u>Video S7</u>) (Fig. S1C). Collectively, *lmn-1, zyg-12* and *sun-1* led to centrosome detachment from the nuclear envelope and an increase in centrosome-cen-trosome distance, supporting a role for the nuclear lamina in promoting centrosome-nuclear envelope cohesion during centrosome migration.

**Figure 2.**
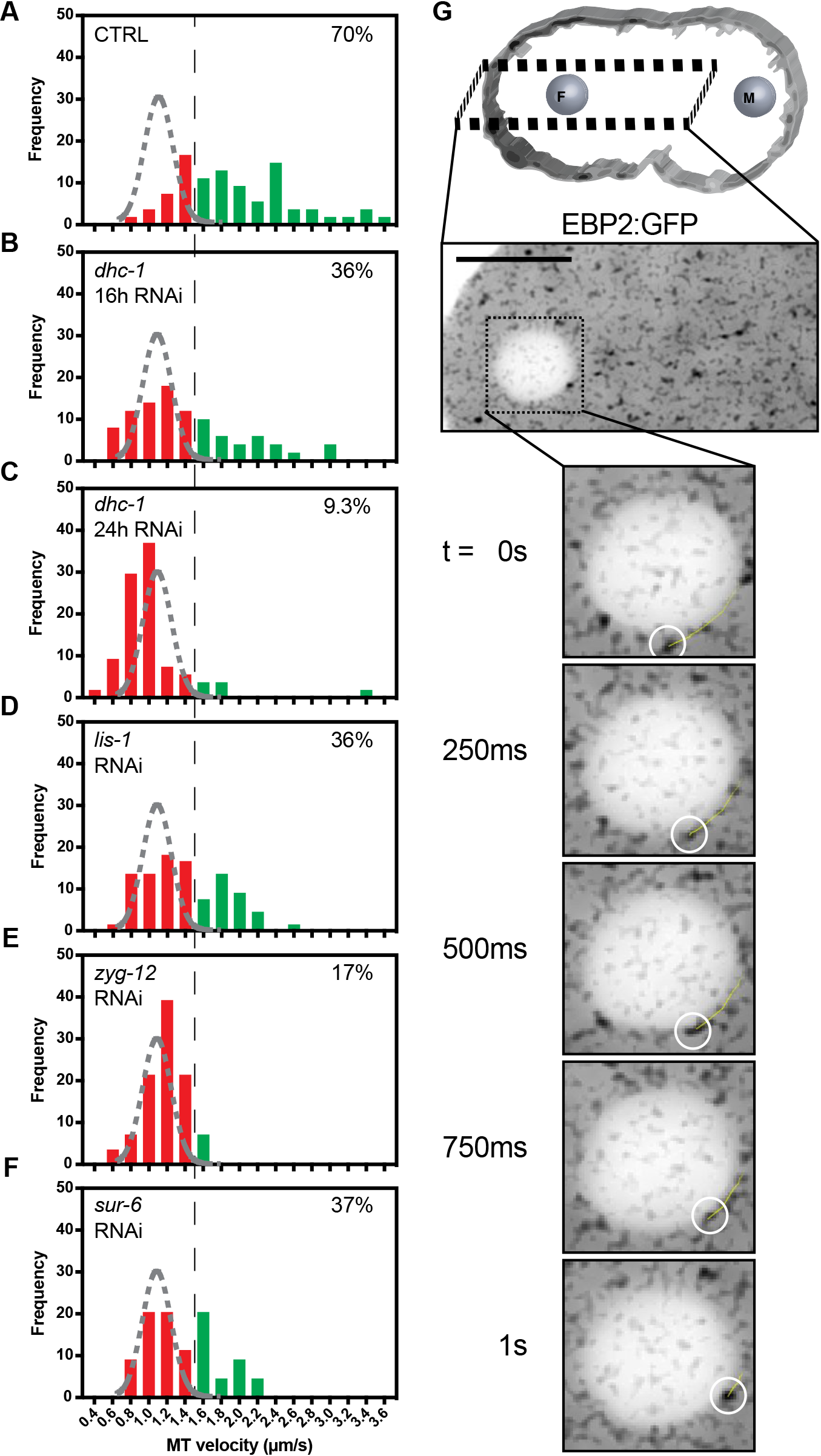
Regulation of Dynein-dependent microtubule growth velocity acceleration at the outer nuclear envelope. (A)(B)(C)(D)(E)(F) Frequency distributions of microtubule velocities for microtubules contacting the nuclear envelope of the female pronucleus prior to pronuclear migration. The black dashed line represents a threshold over which microtubules are assumed to be accelerated by nuclear envelope-tethered Dynein. The grey dashed curve represents a gaussian distribution fit to control cytoplasmic microtubule growth velocities. n of microtubules analyzed per condition is > 50 across > 3 embryos. Percentage inlays represent the proportion of microtubules growing at velocities above the 1.5 μm/s threshold in each condition. (G) Representative single plane confocal image of embryos expressing EBP2:GFP. The yellow trajectory line represents a track for up to 10 frames (2.5 s) following the displayed frame using TrackMate. Scale bar = 10 μm.

### Nuclear envelope-based Dynein motor activity is regulated by PP2A-B55/SUR-6 and ZYG-12

How biochemical regulators such as PP2A-B55/ SUR-6 and components of the LINC complex maintain centrosome positioning in the embryo was unclear. To gain mechanistic insight into centrosome positioning regulation, nuclear envelope-based Dynein motor capability was measured using a previously described approach (Srayko et al., 2005). *C. elegans* embryos expressing the microtubule plus-end binding protein EBP2:GFP were imaged prior to pronuclear migration and microtubule growth velocities were measured (<u>Video S8</u>). Importantly, regulation of global microtubule polymerization rates in RNAi-compromised conditions was measured by tracking cytoplasmic microtubules and was found to be unperturbed (Fig S2). Based on cytoplasmic microtubule growth velocities, we estimated that microtubules with measured mean velocities above 1.5 μm/s are likely to be accelerated by external forces as few microtubules exhibited polymerization rates >1.5 μm/s in the cytoplasm.

Measuring microtubule growth velocities for microtubules interacting with the nuclear envelope on the female pronucleus revealed that >70% of microtubules were accelerated relative to cytoplasmic microtubules (Fig 2A). In control embryos, median cytoplasmic microtubule growth velocity was 1.09 μm/s, or ~0.4 μm/s slower than nuclear associated microtubules (Fig S2A). Reconstituted mammalian Dynein complexes have been found to exhibit average velocities of just below 0.6 μm/s in vitro (Gutierrez et al., 2017). Therefore, our observed 0.4 μm/s average increase in apparent microtubule growth velocities in the vicinity of the nuclear envelope are within Dynein’s ability to accelerate microtubules. Consistent with previous reports, microtubule acceleration in the vicinity of the nuclear envelope was dependent on Dynein, as depletion of *dhc-1* by RNAi led to a reduction in the population of accelerated microtubules in a titratable manner between 16 hour (Fig 2B) and 24 hour (Fig 2C) RNAi treatments. Depleting the Dynein regulator *lis-1* also led to a reduction in the fraction of accelerated microtubules, supporting Dynein’s nuclear envelope-based motoring activity (Fig 2D).

**Figure 3.**
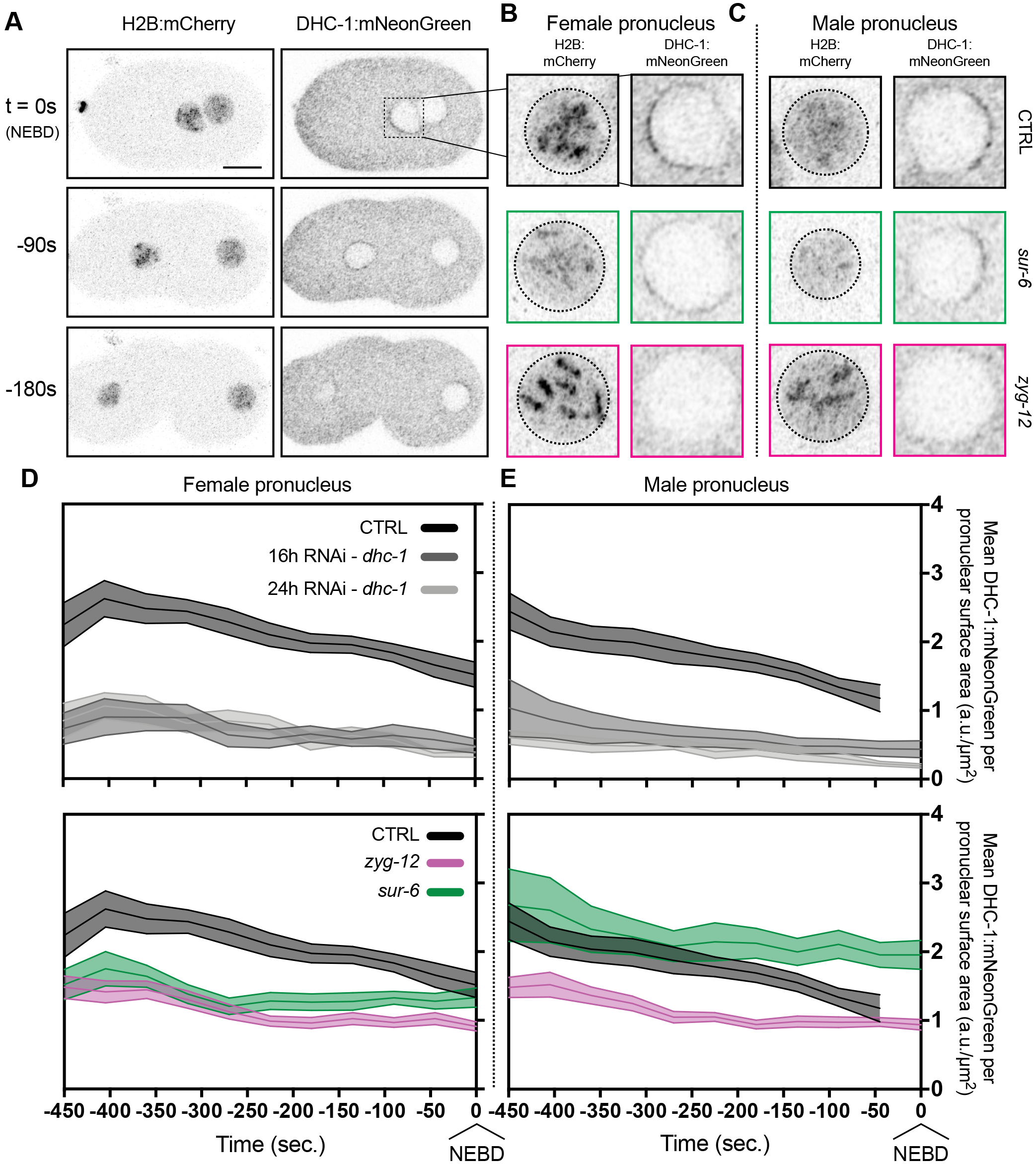
Quantification of endogenous DHC-1:mNeonGreen reveals nuclear envelope localization regulation by PP2A-B55/SUR-6 and ZYG-12. (A) Representative confocal images of embryos expressing mCherry:HIS-58 and endogenous DHC-1:mNeonGreen. mCherry:HIS-58 images represent maximum intensity projections of confocal stacks and DHC-1:mNeonGreen images are single planes. Scale bar = 10 μm. (B)(C) Inlays of (B) female pronuclei and (C) male pronuclei one frame prior to NEBD. (D)(E) Density measurements of DHC-1:mNeonGreen (arbitrary units – a.u.) per surface area of nuclear envelope (μm^2^) on (D) female pronuclei and (E) male pronuclei. n of embryos analyzed per condition is > 14. Time t = 0 s corresponds to NEBD.

**Figure 4.**
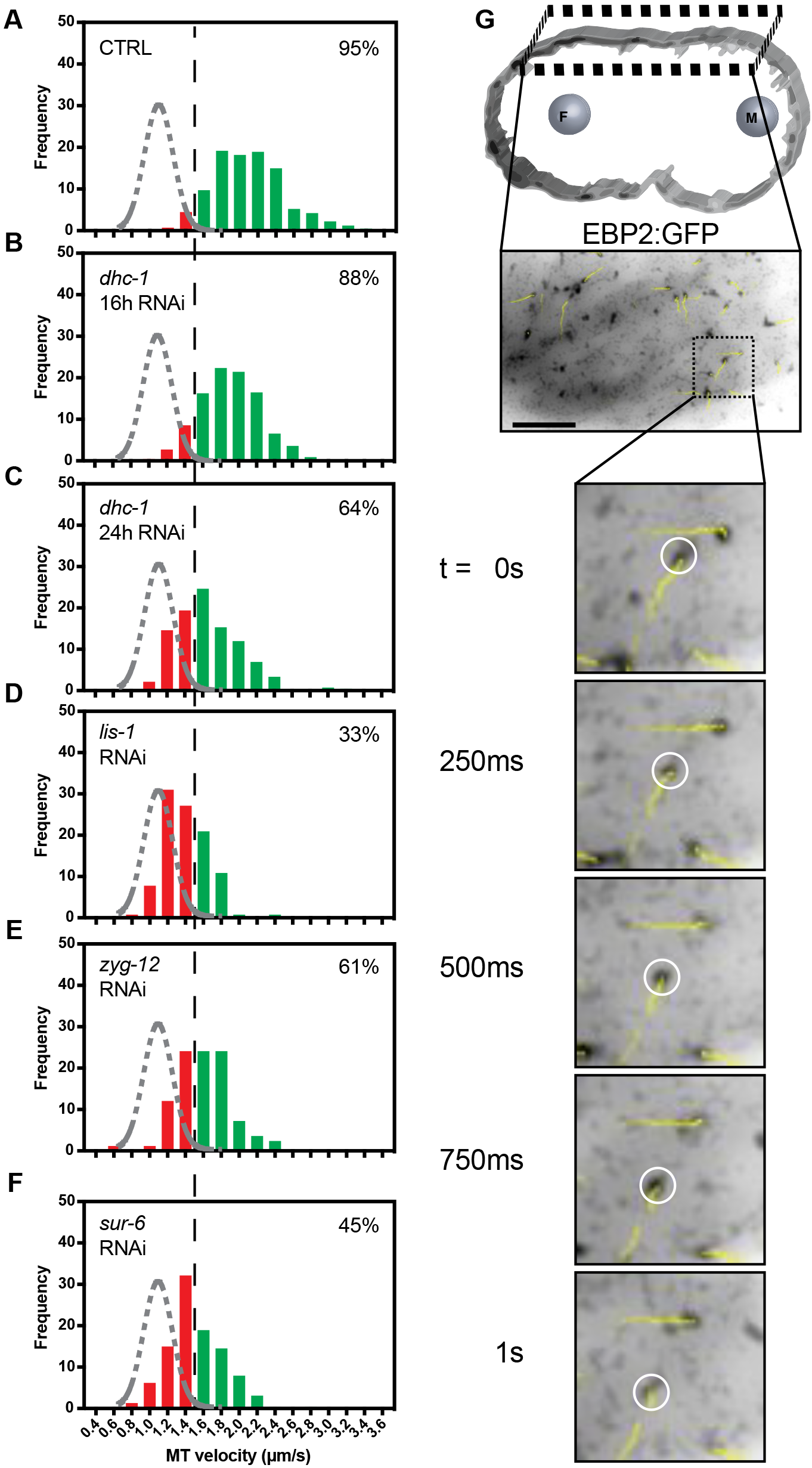
Regulation of Dynein-dependent microtubule growth velocity acceleration at the cortex. (A)(B)(C)(D)(E)(F) Frequency distributions of microtubule velocities for microtubules polymerizing along the cortex of the embryo prior to pronuclear migration as visualized by TIRF microscopy. The black dashed line represents a threshold over which microtubules are assumed to be accelerated by cortical Dynein. The grey dashed curve represents a gaussian distribution fit to control cytoplasmic microtubule growth velocities. n of microtubules analyzed per condition is > 80 across > 3 embryos. Percentage inlays represent the proportion of microtubules growing at velocities above the 1.5 μm/s threshold in each condition. (G) Representative single plane confocal image of embryos expressing EBP2:GFP. Yellow trajectory lines represent tracks for up to 10 frames (2.5 s) following the displayed frame using TrackMate. Scale bar = 10 μm.

To determine how components known to affect centrosome positioning may affect Dynein-mediated pulling forces, microtubule growth velocities of microtubules interacting with the nuclear envelope were measured in *zyg-12* (Fig 2E) and *sur-6* (Fig 2F) depleted embryos. Depleting *zyg-12*, which has previously been shown to be required for DHC-1 localization to the nuclear envelope in the germline and in embryos (Malone et al., 2003), decreased the proportion of accelerated microtubules (Fig 2E). Depleting *sur-6* also led to a decrease in the proportion of accelerated microtubules at the female pronucleus (Fig 2F), suggesting PP2A-B55/SUR-6 may regulate centrosome positioning through microtubule-based Dynein motoring activity.

### PP2A-B55/SUR-6 and ZYG-12 regulate nuclear envelope-based Dynein density

Biochemical and biophysical regulation of centrosome positioning through Dynein could occur through several mechanisms including Dynein affinity for the nuclear envelope (mean amount), Dynein density on the nuclear envelope (amount per surface area) or Dynein motoring activity itself. Based on the above results, we asked how nuclear envelope-based Dynein motor capability could be regulated by PP2A-B55/SUR-6. To distinguish between potential Dynein regulatory mechanisms, we measured amounts of endogenous DHC-1 (Heppert et al., 2018) during mitotic entry on the nuclear envelope (<u>Video S9</u>).

Depletion of *zyg-12* in the embryo caused a significant loss in DHC-1:mNeonGreen density prior to nuclear envelope breakdown on both the female and male pronuclei (Fig 3D, E). ZYG-12-dependent localization of DHC-1 is reflected both in DHC-1 density on the surface of the nuclear envelope as well as in the average amount of DHC-1 on the envelope (Fig S3B).

Depletion of *sur-6* led to insignificant differences in average amounts of DHC-1:mNeonGreen on female and male pronuclei (Fig S3B). However, density measurements of DHC-1 with respect to nuclear envelope surface area revealed female pronuclei had significantly less DHC-1 per unit area compared to their control counterparts, whereas male pronuclei had higher DHC-1 densities (Fig 3D, E). Interestingly, depleting *sur-6* led to significant pronuclear size changes in which female pronuclei were 66% larger and male pronuclei were 18% smaller in volume (Fig S4). Taken together, measuring DHC-1 amounts on pronuclei revealed ZYG-12 regulates Dynein motor activity throughout mitotic entry by regulating DHC-1 localization on the outer nuclear envelope, whereas SUR-6 affects Dynein motor activity, possibly indirectly, through the regulation of pronuclear size, affecting pronuclear DHC-1 densities.

### PP2A-B55/SUR-6 regulates cortical Dynein motoring activity

In addition to nuclear envelope-based Dynein, cortex-associated Dynein has also been shown to be critical for centrosome positioning in the *C. elegans* embryo (De Simone et al., 2016). Microtubules interacting with Dynein tethered to the cortex have been shown to be accelerated in a Dynein-dependent manner during metaphase (Gusnowski and Srayko, 2011). To determine if PP2A-B55/SUR-6 could be regulating centrosome positioning through the regulation of cortical Dynein in addition to nuclear envelope-associated Dynein, untethered microtubules growing in the vicinity of the cortex were imaged using TIRF microscopy (<u>Video S10</u>).

At the cortex, most microtubules grew at rates above the 1.5 μm/s threshold (Fig 4A), indicating prevalent microtubule accelerating forces. The measured polymerization rates were dependent on Dynein in a titratable manner as expected (Fig 4B and 4C), as well as on the Dynein regulator LIS-1 (Fig 4D). Depleting *sur-6* via RNAi led to a reduction in over 50% of the proportion of microtubules being accelerated by Dynein (Fig 4E), suggesting that PP2A-B55/SUR-6 regulates cortical Dynein in addition to nuclear envelope-tethered Dynein. Interestingly, depleting *zyg-12* led to a reduction in the proportion of Dynein-accelerated microtubules (Fig 4E) to the same extent as a 24 hour *dhc-1* depletion (Fig 4C). How ZYG-12 could be regulating cortical Dynein activity is unclear and should be the subject of future work.

### Computational simulations of centrosome migration partially recapitulate PP2A-B55/SUR-6’s role in regulating centrosome positioning through pronuclear size and nuclear envelope Dynein density

Given the pleotropic nature of PP2A-B55/SUR-6-mediated Dynein regulation, we turned to computational simulations to identify phenotypes relevant to centrosome positioning. As a mitotic master regulator, PP2A-B55/SUR-6 is known to regulate several aspects of mitotic entry and exit including mitotic entry timing (Mochida et al., 2009), chromosome decondensation and nuclear envelope reformation (Afonso et al., 2014), as well as cytokinesis initiation (Cundell et al., 2013). To determine if the uncovered roles for PP2A-B55/SUR-6 in affecting pronuclear size (Fig S4), nuclear envelope Dynein density (Fig 3B-E) and cortical Dynein activity (Fig 4F) are sufficient to affect centrosome positioning, a previously described Cytosim-based computational model was used (<u>Video S11</u>) (Fig 5A) (De Simone et al., 2016).

Separating *sur-6* effects, we simulated differential pronuclear sizes, Dynein densities on pronuclear envelopes and cortical Dynein motor activity during centrosome separation. First, pronuclear sizes were modified as to reflect *sur-6* effects while maintaining Dynein densities. Female pronuclei were simulated to have initial volumes 66% larger (μm^3^) than control while male pronuclei were 18% smaller, as measured experimentally. Interestingly, modifying only pronuclear volumes had a subtle yet significant effect on centrosome positioning (Fig 5C). Second, pronuclear Dynein densities were modified to reflect experimental values independently of pronuclear size differences. Decreasing Dynein surface density on the female pronucleus by 25% (a.u./μm^2^) while increasing Dynein surface density on the male pronucleus by 40% led to significant differences in centrosome separation (Fig 5D). Third, cortical Dynein motoring activity was simulated as 50% slower than control to reflect experimental values. Cortical Dynein on its own did not have significant effects on centrosome positioning (Fig S5), or synergistic effects when combined with pronuclear size and Dynein density perturbations (data not shown), suggesting PP2A-B55/SUR-6 regulation of cortical Dynein is not a significant centrosome separation regulatory mechanism.

To determine whether these perturbations collaborate in ensuring centrosome positioning, simulating pronuclear size and Dynein density differences between nuclei simultaneously generated an additive phenotype in which centrosome-centrosome distance was reduced further (Fig 5E). Taken together, the simulations uncovered nuclear envelope-based *sur-6* phenotypes on centrosome positioning reduced centrosome-centrosome distance. These data support a role for PP2A-B55/SUR-6 in regulating centrosome positioning through nuclear size regulation and its effect on Dynein density on the nuclear envelope.

Evidently, the *sur-6* phenotype could only be partially recapitulated through its regulation of nuclear size and nuclear envelope Dynein densities, indicating that *sur-6* likely induces other processes relevant to centrosome positioning that have yet to be uncovered. Another clear centrosome-based *sur-6* phenotype was the initial position of centrosome pairs prior to centrosome separation (Fig 1B). In *sur-6* depletions, centrosome pairs were positioned internally to the embryo as opposed to being proximal to the cortex, potentially disrupting the orientation of microtubule pulling or pushing forces against the cortex. Simulating centrosome migration in embryos in which centrosomes were internal to the embryo (Fig 5F) abolished centrosome separation, suggesting initial centrosome positioning with respect to the cortex is required for centrosome migration (Fig 5G).

### Compromising Dynein-mediated pulling forces and centro-some-nuclear envelope cohesion simultaneously abolishes centrosome migration

To determine how PP2A-B55/SUR-6 regulation of centrosome positioning is coordinated with LINC complex-mediated centrosome-nuclear envelope tethering, we depleted components of both pathways simultaneously via RNAi. As described previously, DHC-1 outer nuclear envelope localization is dependent on the hook protein ZYG-12 (Malone et al., 2003 and Fig 3D, E) and both DHC-1 and ZYG-12 are required for centrosome-nuclear envelope cohesion (Fig 1). Here, we titrated the amount of DHC-1 required for centrosome-nuclear envelope cohesion while affecting centrosome separation (Fig 1C) and motoring ability at the nuclear envelope (Fig 2B) and at the cortex (Fig 4B). Compromising *lis-1* via partial RNAi also provided a condition in which centrosomes remained tethered to nuclear envelopes (Fig S1B) while compromising Dynein motoring ability at the nuclear envelope (Fig 2D) and at the cortex (Fig 4D). Partial RNAi depletions of *dhc-1* and *lis-1* therefore allow us to test the contribution of Dynein-mediated pulling forces in addition to centrosome-nuclear envelope tethering in positioning centrosomes in the *C. elegans* embryo.

Testing if PP2A-B55/SUR-6 and the nuclear lamina collaborate in positioning centrosomes, *sur-6* and *lmn-1* were co-depleted in embryos (<u>Video S12</u>). Centrosomes migrated to the center of embryos detached from the male pronuclear envelope and with compromised centrosome-centrosome distances (Fig 6B). The measured centrosome positioning phenotype in *sur-6* and *lmn-1* (Fig 6D) was only slightly more severe than in *sur-6* alone (Fig 1B), although a combination of centrosome separation and nuclear envelope tethering phenotypes were observed. Taken together, co-depletion of *sur-6* and *lmn-1* suggests centrosome positioning is achieved through the complimentary regulation of Dynein-me-diated pulling forces and centrosome-nuclear envelope tethering.

**Figure 5.**
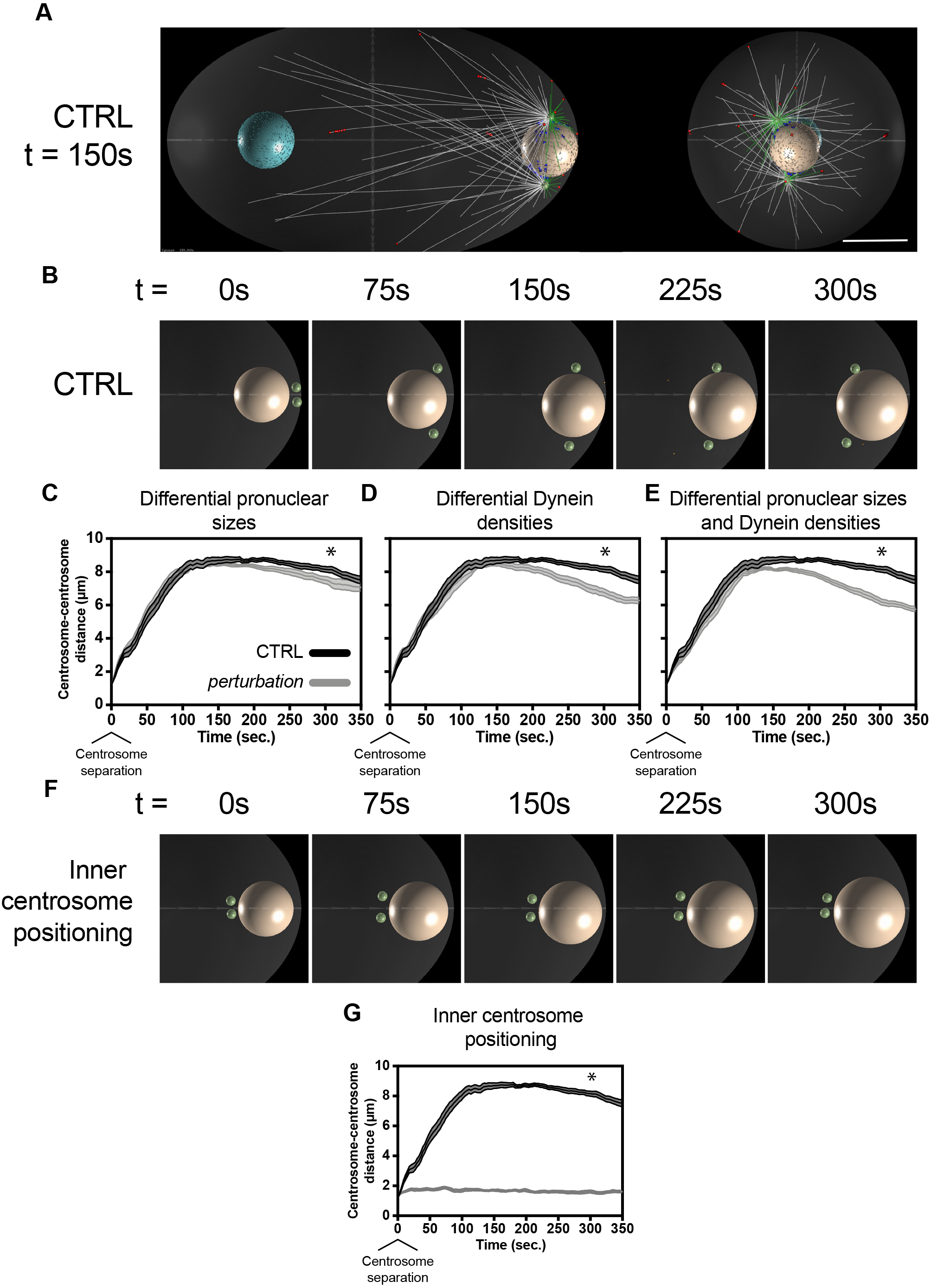
Cytosim-based computational simulations predict differences in pronuclear size and nuclear envelope-tethered Dynein density can affect centrosome positioning, while initial inner centrosome positioning is dominant. (A) Representative image of Cytosim-based centrosome positioning simulations at 150 s post centrosome separation. Red and blue dots represent Dynein-microtubule binding at the cortex and at the nuclear envelope, respectively. White microtubules represent inwardly directed microtubules and green microtubules represent rearward directed microtubules with respect to centrosomes (green). The rightmost image represents an end-on view of the posterior of the embryo. Scale bar = 10 μm. (B) Representative images of embryo inlays through early centrosome migration. Male pronuclei are displayed in beige and centrosomes in green. (C) Predicted centrosome-centrosome distances in control embryos and in embryos with 18% smaller male pronuclei and 66% larger female pronuclei (μm^3^). (D) Predicted centrosome-centrosome distances in control embryos and in embryos with 40% denser Dynein on male pronuclei and 25% sparser Dynein on female pronuclei (a.u./μm^2^). (E) Predicted centrosome-centrosome distances in control embryos and in embryos with both differentially sized pronuclei and Dynein densities as in (C) and (D). (F) Representative images of simulated centrosome separation with inner initial centrosome positioning. (G) Predicted centrosome-centrosome distances in control embryos and in embryos with inner initial centrosome positioning. All simulated perturbations reflect experimental measurements of *sur-6* RNAi phenotypes. Error bars are SEM and are represented as shaded areas around means of 10 simulation runs per condition. Time t = 0 s corresponds to centrosome separation. * = p < 0.05 at 300 s post centrosome separation.

**Figure 6.**
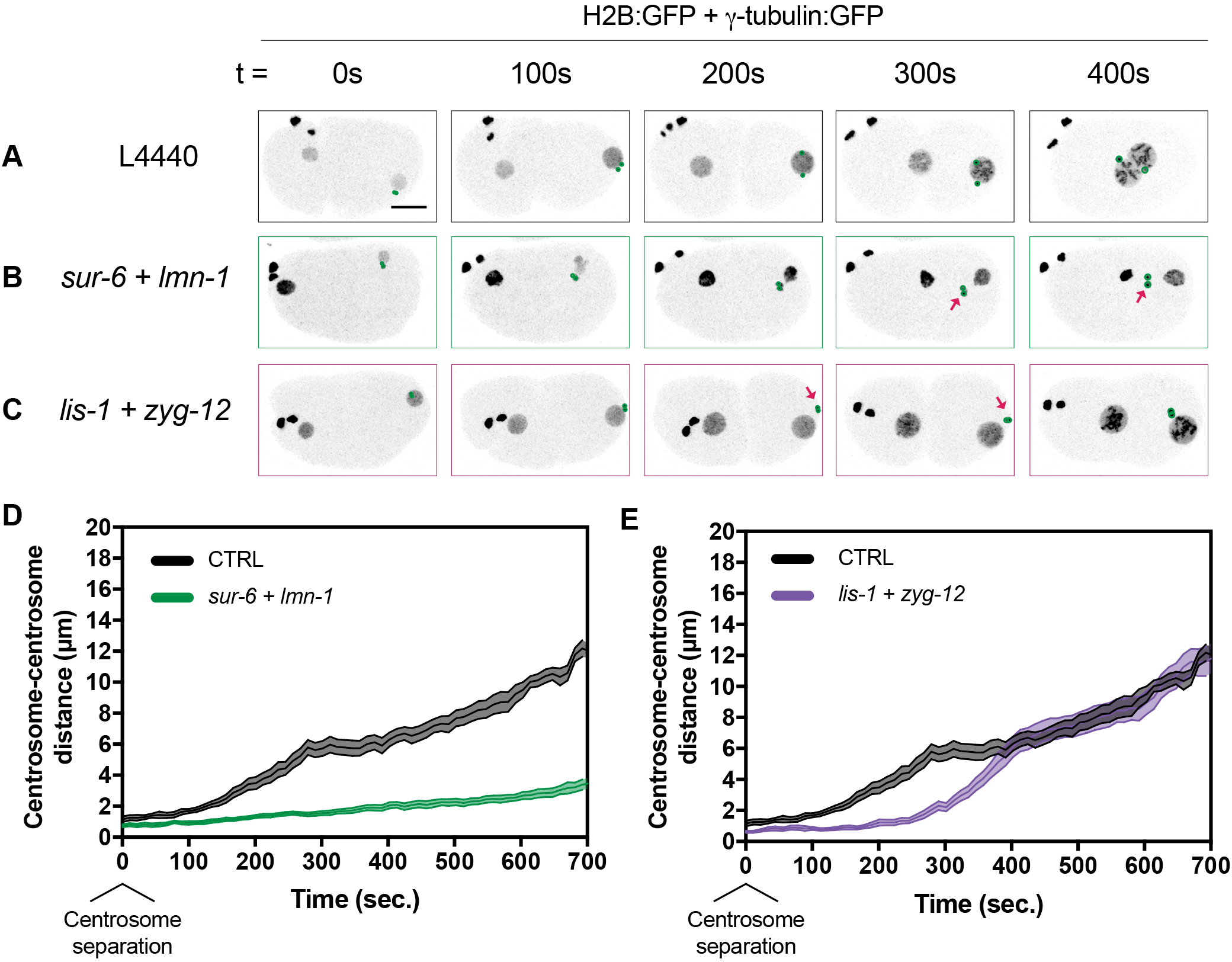
Co-depleting *sur-6* and *lmn-1* or *lis-1* and *zyg-12* leads to centrosome attachment and centrosome separation defects. (A)(B)(C) Representative maximum intensity projections of embryonic confocal stacks. All RNAi treatments were performed with L4 stage worms expressing H2B:GFP and γ-tubulin:GFP for 24 hours via feeding unless otherwise noted. Circles highlight individual centrosomes and arrows denote detached pairs of centrosomes. Scale bar = 10 μm. (D)(E) Centrosome distance was measured in 2D using maximum intensity projections of confocal stacks. Error bars are SEM and are represented as shaded areas around means. Time t = 0 s corresponds to centrosome separation.

To gain mechanistic insight, we then tested whether Dynein-based pulling forces collaborate with centrosome-nuclear envelope tethering to position centrosomes separately from *sur-6* effects on pronuclear size and initial centrosome positioning. We reasoned that simultaneously compromising other components of the Dynein motoring machinery and of the LINC complex would equally abrogate centrosome separation, leading to a combination of centrosome positioning phenotypes similar to *sur-6* + *lmn-1.* To test this, we co-depleted *lis-1* and *zyg-12* using partial RNAi depletions (<u>Video S13</u>) (Fig 6C). Centrosomes became both detached from the male pronuclear envelope and centrosome separation was severely impaired (Fig 6E). We conclude that Dynein-mediated pulling forces regulated by LIS-1 and centrosome-nuclear envelope tethering regulated by ZYG-12 collaborate for proper centrosome positioning during the early stage of centrosome separation.

Taken together, *sur-6* and *lmn-1* or *lis-1* and *zyg-12* co-depletions led to additive phenotypes in which centrosomes are detached from the male pronucleus during centrosome migration and fail to separate from one another. In both conditions, centrosome separation is severely compromised, suggesting that Dynein-based pulling forces and cen-trosome-nuclear envelope tethering collaborate during early stages of centrosome separation through parallel pathways.

## DISCUSSION

By combining quantitative live cell-imaging, particle tracking and computational simulations, we determined that PP2A-B55/SUR-6 and the nuclear lamina collaborate in positioning centrosomes through separate pathways. Although the mechanism by which *sur-6* affects nuclear size is unclear, PP2A-B55/ SUR-6 regulation through the nuclear export of its negative regulator during mitotic entry in Drosophila has been described previously (Wang et al., 2013) and may prove to be important in the worm embryo. Importantly, nucleo-cytoplasmic trafficking has been shown to regulate nuclear and chromosome size scaling in *C. elegans* (Ladouceur et al., 2015) as well as nuclear expansion rates (Boudreau et al., 2018), making nucleo-cytoplas-mic trafficking a likely regulatory candidate in ensuring proper PP2A-B55/SUR-6 function. Despite considering several *sur-6* phenotypes, PP2A-B55/SUR-6 is likely to regulate several other components relevant to centrosome migration, including the regulation of centriole duplication, as has been described for mitotic exit leading to the embryonic two cell stage (Song et al., 2011).

The respective roles of nuclear envelope and cortical Dynein in pulling on centrosomes have recently become clearer (De Simone et al., 2016, 2017, 2018). How Dynein-based pulling forces are transmitted through the LINC complex to allow for centrosome separation and pronuclear migration have yet to be uncovered. Compromising LINC complex components *zyg-12* or *sun-1* caused similar centrosome separation phenotypes, which were also phenocopied by compromising the nuclear lamina as a whole. Although LMN-1 and SUN-1 are likely required for Dynein localization on the nuclear envelope in addition to ZYG-12, models in which the LINC complex functions in tethering centrosomes to the nuclear envelope and in promoting Dynein motoring activity on the nuclear envelope surface are not mutually exclusive. Partial depletions of *dhc-1* and *lis-1* led to reduced Dynein motor capacity at the nuclear envelope (Fig 2B, D) and at the cortex (Fig 4B, D), while centrosome-nuclear envelope cohesion was maintained (Fig 1C and Fig S1). In fact, quantification of endogenous nuclear envelope-associated DHC-1 suggests that in 16 hour *dhc-1* depletions, DHC-1 levels are lower on the nuclear envelope compared to *zyg-12* embryos (Fig 3). This indicates that centrosomes are tethered to the nuclear envelope through Dynein as well as independently of Dynein. Centrosome-nuclear envelope tethering has also been suggested to occur through ZYG-12 dimerization between outer nuclear envelope and centrosome-based pools of the hook protein (Malone et al., 2003). Alternative tethering mechanisms include ZYG-12-microtubule binding, supported by the presence of microtubule binding domains in most mammalian hook proteins (Walenta et al., 2001), as well through nuclear pore-microtubule binding, as identified in several metazoans (reviewed in Goldberg, 2017).

Using computational simulations, several of PP2A-B55/ SUR-6’s uncovered roles were evaluated separately (Fig 5). Perturbing pronuclear size (Fig 5C) and nuclear envelope-based Dynein density (Fig 5D) led to defects late in centrosome positioning, potentially indicating defects in centrosome centration rather than centrosome separation. Similarly, defects in initial centrosome positioning relative to the cortex (Fig 5F, G) reveal early centrosome separation requirements consistent with previous observations in the initial positioning of the male pronucleus (De Simone et al., 2017). Although the molecular mechanisms by which PP2A-B55/SUR-6 may regulate initial centrosome positioning remain elusive, elevated Dynein densities on the male pronucleus in *sur-6* (Fig 3E) may contribute to premature centrosome migration towards the interior of the embryo. Interestingly, centrosome distance in *lis-1* + *zyg-12* recovered approximately 400 s following centrosome separation (Fig 6E), indicating that a collaboration between Dynein-based pulling forces and cen-trosome-nuclear envelope cohesion may only be required early during centrosome separation. On the other hand, initial centrosome positioning in sur-6 (Fig 1B) may play a dominant role beyond centrosome separation (Fig 1F), revealing potential roles of cortical Dynein or other regulatory mechanisms at later stages that could not be captured by our computational simulations. Importantly, despite centrosome distance recovery in *lis-1* + *zyg-12*, severe chromosome congression and segregation phenotypes were prevalent (Video S12), indicating that early centrosome separation and migration are likely critical for subsequent mitotic events.

In conclusion, we propose a model through which Dynein-mediated pulling forces and centrosome-nuclear envelope cohesion are coordinated during mitotic entry to ensure proper centrosome migration and positioning. The collaboration between these pathways broadly and between PP2A-B55/SUR-6 and Lamin/LMN-1 specifically are critical for ensuring mitotic fidelity and are likely to be utilized across tissues and organisms.

## METHODS

### Computational simulations

Computational modeling of centrosome dynamics was performed using Cytosim under parameters described previously (De Simone et al., 2016). Brownian motion of elastic fibers and solids in a constant homogenous viscous medium are described by overdamped Langevin equations. The embryo is simulated as an ellipsoid (50 × 30 × 30 μm), confining all embryonic elements. Centrosomes serve as microtubule nucleation sites. Microtubules undergo dynamic instability and can interact with dynein motors, evenly distributed on pronuclear surfaces and the cell cortex. The female and male pronuclei are at their respective anterior and posterior locations of the embryo at t = 0. Centrosomes are 1.2 μm apart and are located between the male pronucleus and the posterior cortex.

Each condition was simulated 10 times. The position of centrosomes (solids) was exported from Cytosim and centrosome distance was measured using a script written in MATLAB (Math-Works). All changesin simulation parameters, reflectingdifferent perturbations, were performed using experimentally obtained values.

### *C. elegans* use, RNAi and microscopy

Worm strains were grown and maintained at 20°C using standard procedures. Bacterial strains containing a vector expressing dsRNA under the IPTG promoter were obtained from the Ahringer library (from Bob Goldstein’s laboratory, University of North Carolina at Chapel Hill, Chapel Hill, NC, USA). Targets were confirmed by sequencing.

Worm strains used were TH32 (pie-1::bg-1::GFP + unc-119(+), pie-1::GFP::H2B + unc-119(+)) and MAS37 (pie-1p::ebp-2::GFP +unc-119(+)). DHC-1:mNeonGreen quantification experiments were performed in a DHC-1:mNeon-Green and mCherry:HIS-58 strain made by crossing LP560 (cp268[dhc::mNG-C1^3xFlag]) I. with [pie-1p::mCherry::his-58 + unc-119(+)] IV. obtained through a backcross from OD426.

For centrosome distance and DHC-1:mNeonGreen density measurements, worm embryos were mounted in Egg buffer (118 mmol/L NaCl, 48 mmol/L KCl, 2 mmol/L CaCl_2_, 2 mmol/L MgCl_2_, 25 mmol/L HEPES, pH 7.3) between a 1.5 coverslip and a microscope slide spaced by 22.81μm glass beads (Whitehouse Scientific) and sealed with Valap (1:1:1 lanolin, petroleum jelly, and parafilm wax). Embryos were then imaged on a Nikon A1r resonant scanning confocal microscope using a ×60 Apo water immersion objective (Nikon), GaASP PMT detectors and NIS-Elements (Nikon) at 22°C.

Cytoplasmic and cortical microtubule polymerization dynamics were measured using embryos mounted in Egg buffer between a 1.5 coverslip and a 4% agarose pad in Egg buffer and a microscope slide, and sealed with Valap. For cytoplasmic microtubule experiments, embryos were then imaged on a Nikon A1r resonant scanning confocal microscope using the Galvano scanner, a ×60 Apo TIRF oil immersion objective (Nikon), GaASP PMT detectors and NIS-Elements (Nikon) at 22°C. For cortical microtubule experiments, embryos were then imaged on a Nikon TIRF microscope using a ×100 Apo TIRF oil immersion objective (Nikon), an Andor iXon3 EMCCD camera and NIS-Elements (Nikon) at 22°C.

### Image analysis

All image analysis was performed using FIJI (Schindelin et al., 2012). For centrosome tracking, Trackmate (Tinevez et al., 2017) was used with a DOG detection and the Simple LAP tracker. For microtubule plus-tip tracking, Track-mate was used with a DOG detection and the Linear motion LAP tracker. Pronuclear size, DHC-1:mNeonGreen fluorescence intensity measurements and image processing was performed using custom FIJI plugins available upon request.

## SUPPLEMENTAL INFORMATION

Supplemental Information includes five figures and can be found with this article. Videos are hyperlinked within the article’s text and are available here: https://github.com/viboud12/Centrosome-positioning-bioRxiv.

## ACKNOWLEDGMENTS

We greatly thank Jennifer Heppert for reagents and Tony Perdue, Director of UNC Chapel Hill’s department of Biology microscopy core for support. We would like to thank Kevin Cannon, Carlos Patiño-Descovich and Michael Werner for critical reading of the manuscript. V.B. was supported by predoctoral fellowships from the *Fonds de Recherche Santé-Québec* (FRQS). This study was also supported by National Science Foundation CAREER Award 1652512 to P.S.M.

## CONTRIBUTIONS

V.B., R.C., and A.E. designed and carried out experiments in *C. elegans* embryos and performed image and data analysis. M.S. performed computational simulations. V.B. wrote FIJI-based image analysis code and performed image analysis. V.B. and P.S.M. wrote the manuscript.

---

**Figure S1.**
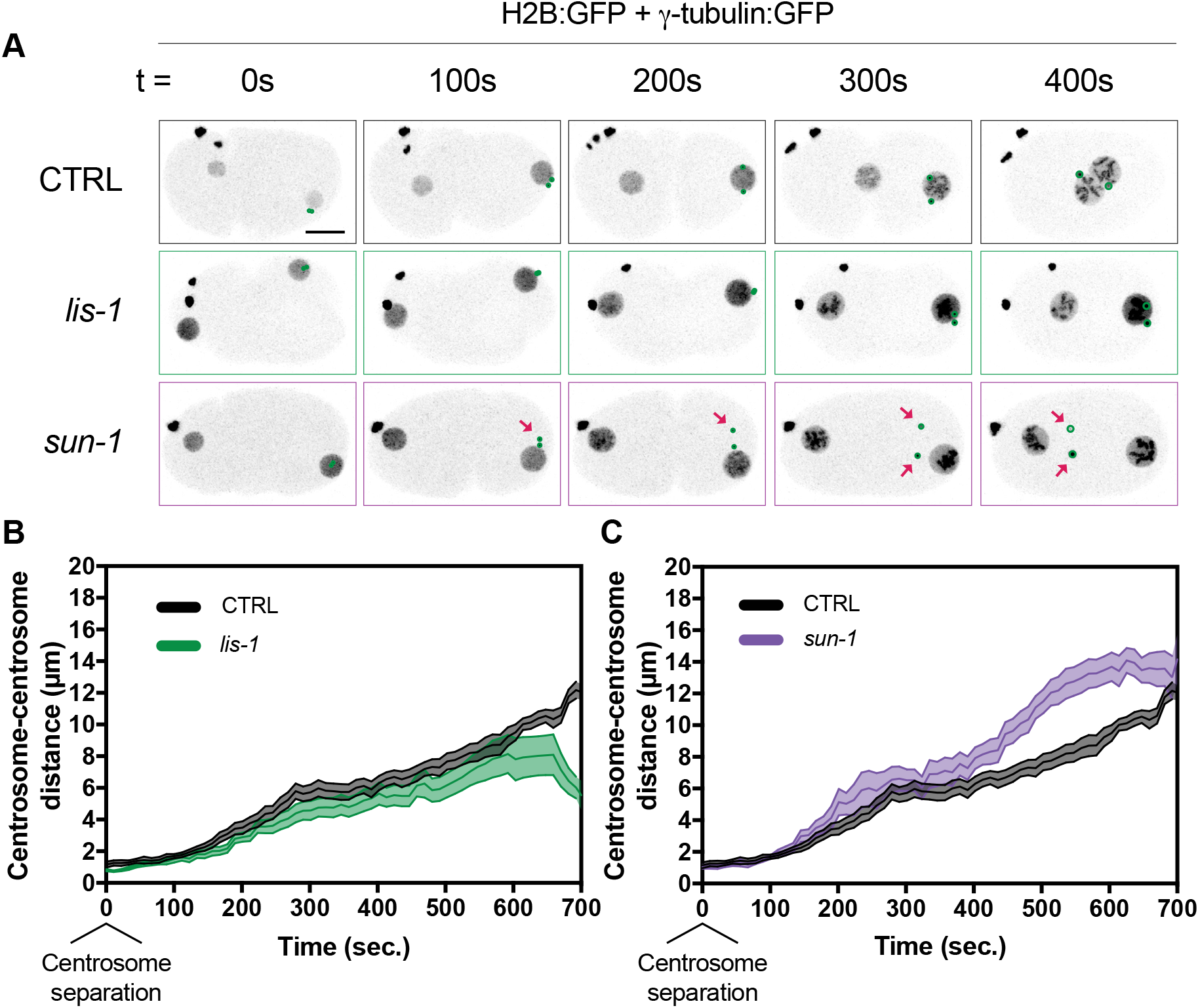
*lis-1* and *sun-1* RNAi affect centrosome separation differentially. (A) Representative maximum intensity projections of embryonic confocal stacks. All RNAi treatments were performed with L4 stage worms expressing H2B:GFP and γ-tubulin:GFP for 24 hours via feeding unless otherwise noted. Scale bar = 10 μm. (B)(C) Centrosome distance was measured in 2D using maximum intensity projections of confocal stacks. Error bars are SEM and are represented as shaded areas around means. n of embryos analyzed per condition > 15. Time t = 0 s corresponds to centrosome separation.

**Figure S2.**
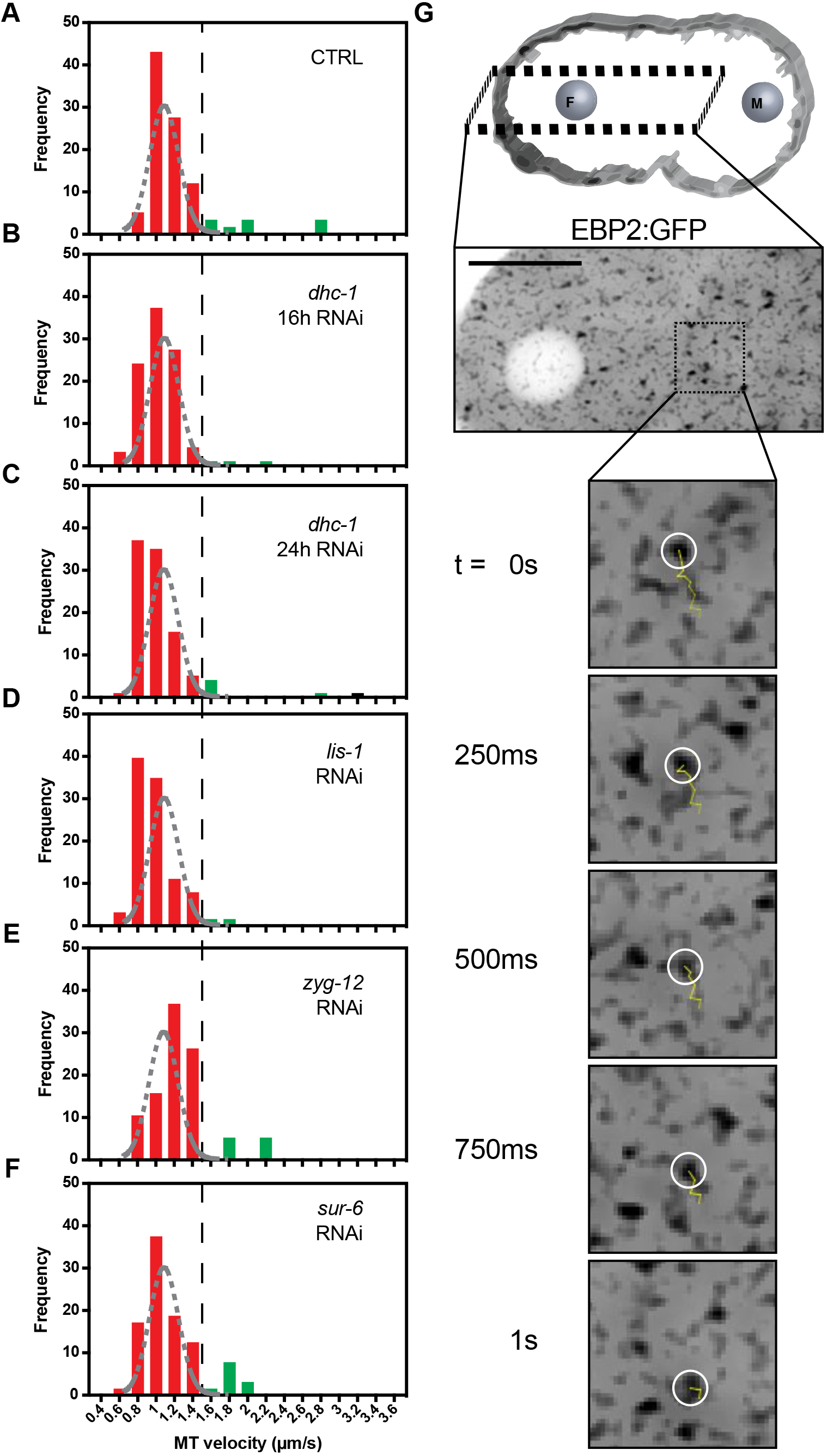
Cytoplasmic microtubule growth velocities prior to pronuclear migration. (A)(B)(C)(D)(E)(F) Frequency distributions of cytoplasmic microtubule growth velocities prior to pronuclear migration. The black dashed line represents a threshold over which microtubules are assumed to be accelerated. The grey dashed curves represent a gaussian distribution fit to control cytoplasmic microtubule growth velocities. n of microtubules analyzed per condition is > 50 across > 3 embryos. Frequency distributions between RNAi conditions were statistically non-significantly different (p > 0.05) using both Mann-Whitney and Kolm-ogorov-Smirnov tests. (G) Representative single plane confocal image of embryos expressing EBP2:GFP. The yellow trajectory line represents a track for up to 10 frames (2.5 s) following the displayed frame using TrackMate. Scale bar = 10 μm.

**Figure S3.**
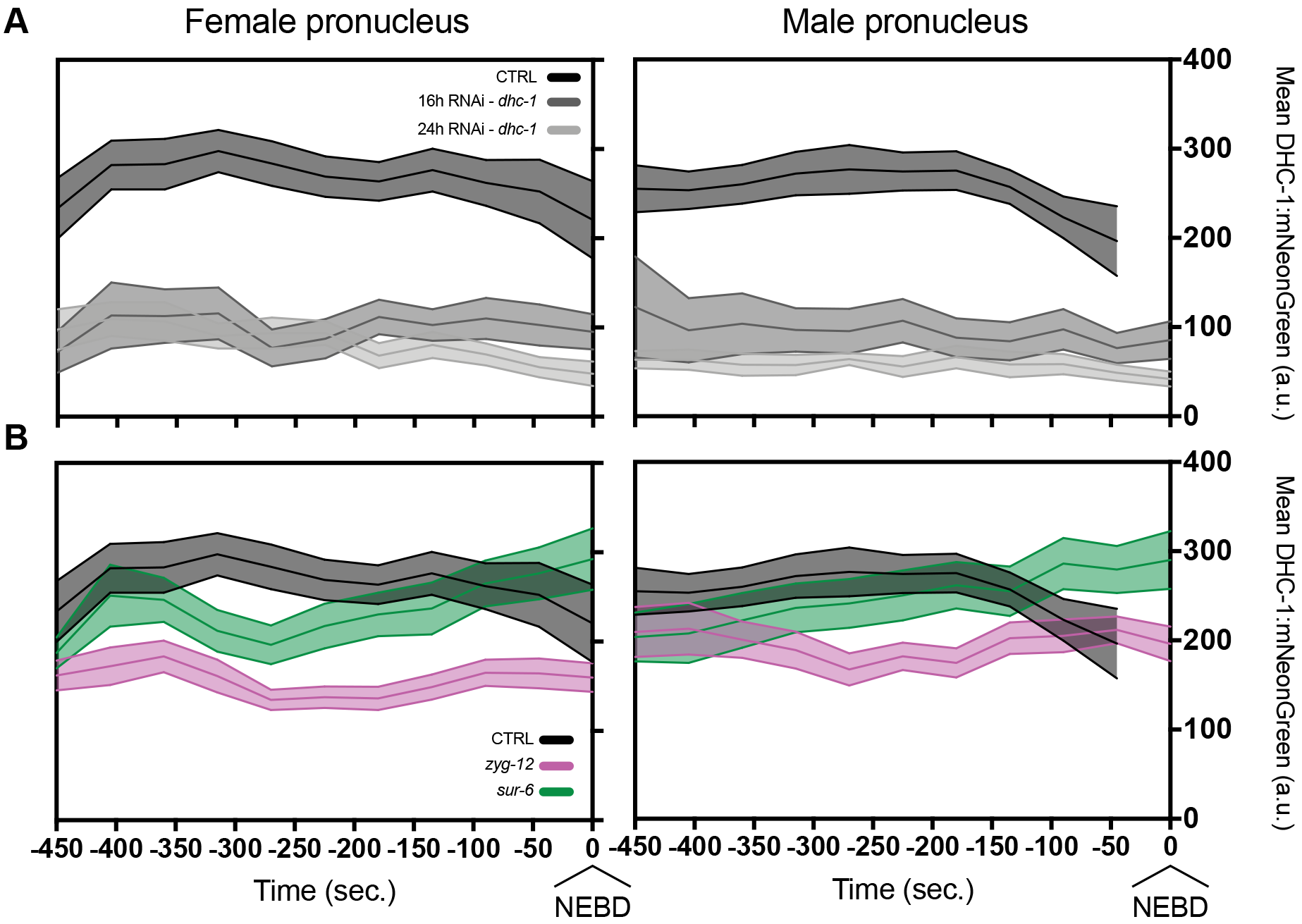
Quantification of endogenous DHC-1:mNeonGreen mean amounts on female and male pronuclear envelopes. (A)(B) Mean amounts of DHC-1:mNeonGreen (arbitrary units – a.u.) on the nuclear envelope in (A) 16 hour and 24 hour *dhc-1* RNAi treatments and (B) 24 hour *zyg-12* and *sur-6* RNAi treatments. Quantifications were performed using single slices of confocal stacks from embryos expressing mCherry:HIS-58 and endogenous DHC-1:mNeonGreen. n of embryos analyzed per condition is > 14. Time t = 0 s corresponds to NEBD.

**Figure S4.**
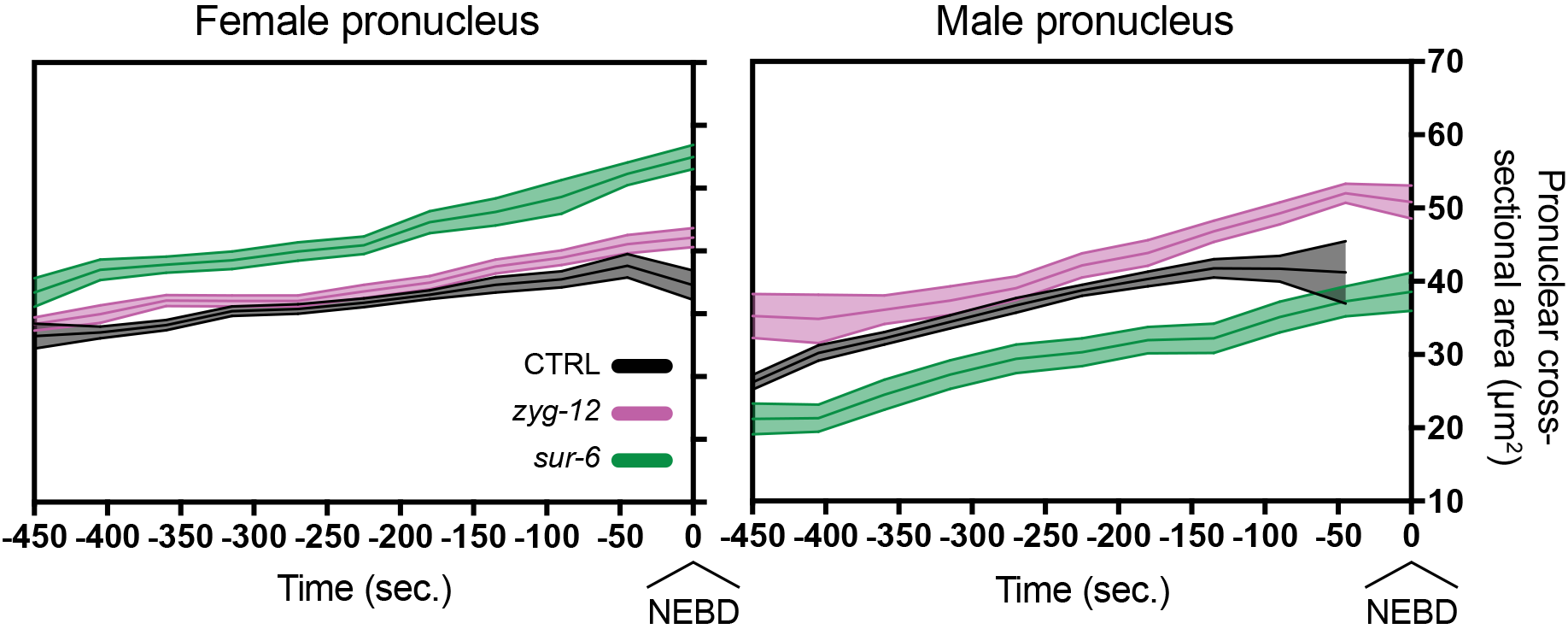
Pronuclear cross-sectional area measurements during pronuclear expansion. Pronuclear cross-sectional areas (μm2) were measured using maximum intensity projections of mCherry:HIS-58 confocal stacks from embryos expressing mCherry:HIS-58 and endogenous DHC-1:mNeonGreen. Female and male pronuclei were analyzed in 24 hour *zyg-12* and *sur-6* RNAi treatments. n of embryos analyzed per condition is > 14. Time t = 0 s corresponds to NEBD.

**Figure S5.**
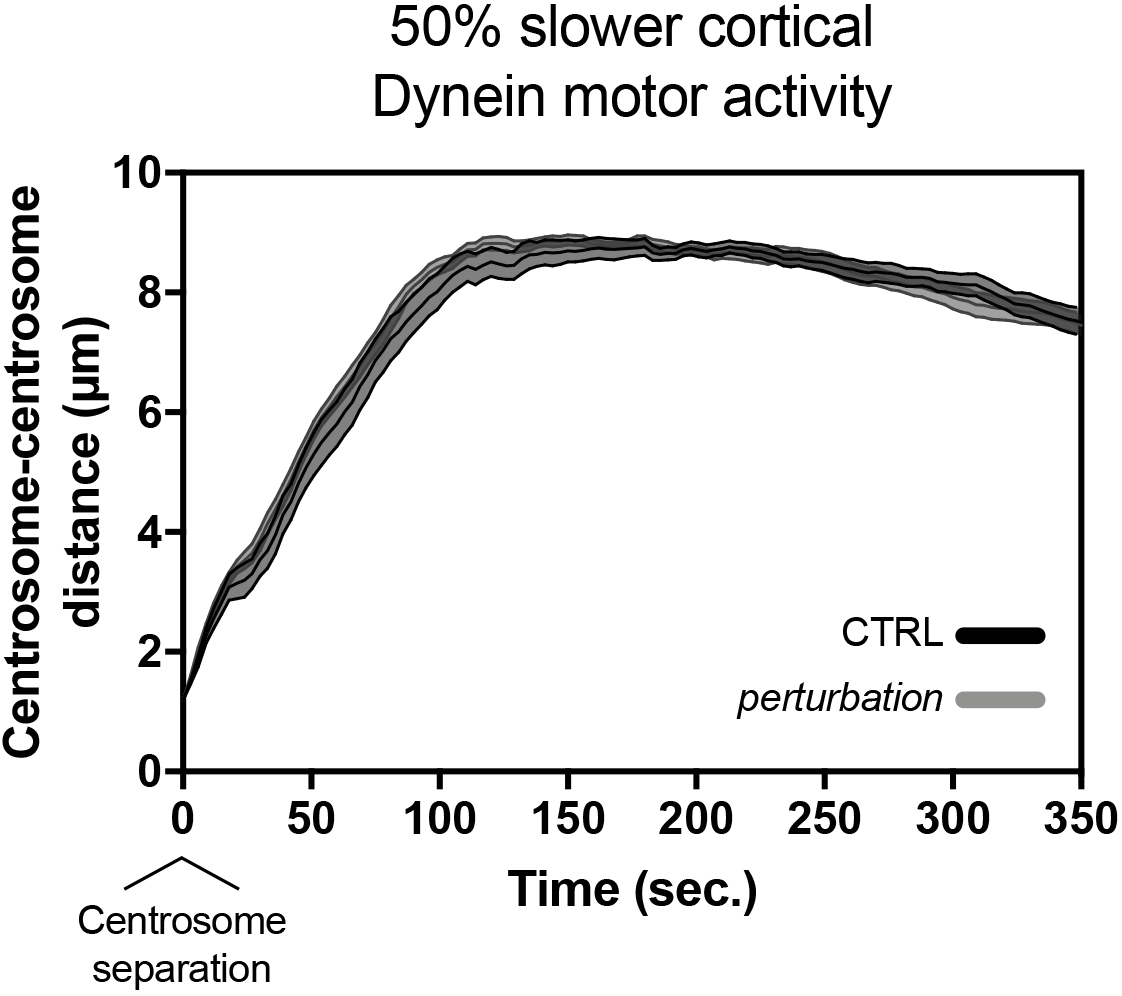
Cytosim-based computational simulations predict 24 hour *sur-6* RNAi effects on cortical Dynein motoring activity are not sufficient to centrosome positioning. Predicted centrosome-centrosome distances in simulated control embryos and in embryos with a 50% reduction in Dynein motoring velocity. Error bars are SEM and are represented as shaded areas around means of 10 simulation runs per condition. Time t = 0 s corresponds to centrosome separation.

